# Candidate targets of copy number deletion events across 17 cancer types

**DOI:** 10.1101/2022.06.29.498080

**Authors:** Qingyao Huang, Michael Baudis

## Abstract

Genome variation is the direct cause of cancer and driver of its clonal evolution. While the impact of many point mutations can be evaluated through their modification of individual genomic elements, even a single copy number aberration (CNA) may encompass hundreds of genes and therefore pose challenges to untangle potentially complex functional effects. However, consistent, recurring and disease-specific patterns in the genome-wide CNA landscape imply that particular CNA may promote cancer-type-specific characteristics. Discerning essential cancer-promoting alterations from the inherent co-dependency in CNA would improve the understanding of mechanisms of CNA and provide new insights into cancer biology and potential therapeutic targets. Here we implement a model using segmental breakpoints to discover non-random gene coverage by copy number deletion (CND). With a diverse set of cancer types from multiple resources, this model identified common and cancer-type-specific oncogenes and tumor suppressor genes as well as cancer-promoting functional pathways. Confirmed by differential expression analysis of data from corresponding cancer types, the results show that for most cancer types, despite dissimilarity of their CND landscapes, similar canonical pathways are affected. In 25 analyses of 17 cancer types, we have identified 19 to 169 significant genes by copy deletion, including RB1, PTEN and CDKN2A as the most significantly deleted genes among all cancer types. We have also shown a shared dependence on core pathways for cancer progression in different cancers as well as cancer type separation by genome-wide significance scores. While this work provides a reference for gene specific significance in many cancers, it chiefly contributes a general framework to derive genomewide significance and molecular insights in CND profiles with a potential for the analysis of rare cancer types as well as non-coding regions.

## 1. Introduction

Cancer genomes are characterized by a wide range of mutations in comparison to the unaltered germline genome. These “somatic” mutations emerge during an individual’s life time and may accumulate sufficiently to lead to malignant transformation and tumorigenesis. Oncogenic mutations can impact the regulation and level of gene expression as well as the completeness and properties of gene products. While deviation from the physiological state typically impair cell viability, two features inherent to malignant transformation, genome instability and high replication rate, frequently promote the generation of a large pool of somatic genome alterations. This pool potentiates the selection of the sporadic cases where the mutated genome promotes a growth advantage and protection from apoptotic mechanisms. However, since most variations do not confer a strong growth advantage[1], the detection of the few key cancer-promoting variations hidden in a complex mutational landscape constitutes a major challenge in cancer genome research.

Depending on the structure of somatic variations, they can be grouped into small-scale sequence alterations, including single nucleotide variations (SNVs), small insertions and deletions (INDELs), and structural variations, including copy number aberrations (CNAs). While the former affects isolated genetic elements, CNAs change the dosage of the covered genetic elements in the affected segment and also may disrupt the local genomic context, e.g. by affecting regulatory elements.

Point mutations have been reported and extensively studied for their functional impact. Affected genes can be evaluated regarding their relevance for oncogenesis through the general effect of their mutations. Briefly, cancer related genes are subdivided into two functional groups: oncogenes, of which gain-of-function (GOF) mutations promote proliferation or inhibit regulatory mechanisms and tumor suppressor genes (TSGs), of which loss of function (LOF) mutations confer a negative impact on cell cycle control and other cellular surveillance functions [2]. The principal modes of action of oncogenes and TSG exhibit differing mutational characteristics. Namely, mutations for oncogenes tend to recur at the same locus; while mutations for TSGs scatter along the coding sequence (CDS) Accordingly, in Catalogue of Somatic Mutations in Cancer (COSMIC) database [3], mutations are classified with a so-called “20/20 rule”: to classify a gene as an oncogene, 20% of all the mutations within a gene’s CDS recorded in database, need to reside at the same locus.; whereas to classify one as a TSG, 20% of recorded mutations need to be inactivating mutations but they mostly do not overlap in their location [1].

In analogy to the diverging functional attributions of point mutations for oncogenes and TSGs, CNAs can be divided into amplifications and deletions. While deletion of a fraction of gene results in LOF due to truncated or untranscribed gene product, only when the entire CDS and potentially the regulatory regions outside CDS are amplified, a GOF arises.

On the mechanistic level, CN gains and losses emerge from erroneous recombination during DNA replication [4] but present unique processes. Extrachromosomal oncogenes have been detected as copy number gain events [5, 6], while chromothripsis - chromosome shattering and rejoining of clustered segments - has been described as a phenomenon in cancer, which disrupts the genetic elements in the region and can result in CN deletions [7]. On the tumor evolution perspective, CN loss events tend to precede CN gain events, suggesting their different roles in oncogenesis [8]. Taken together, the mechanism and impact for amplification and deletion are dissimilar. In this study, we particularly focus on the copy number deletion (CND) patterns modeling the gene inactivation incorporating its unique feature of introducing segmental breakpoints within a gene’s CDS.

Whereas point mutations target one particular genetic element at the specific location, a single CND potentially affects hundreds of genetic elements with a subsequent co-segregation of the affected genes. In addition, overall CNA involvement is highly correlated with the disease stage [9, 10, 11, 12], indicating an accumulation of unrepaired replication defects instead of a predominant selection of driver events. These factors present additional layers of complexity to distinguish the significant genes within large segmental CNA. Yet, CNAs manifest as genome-wide landscapes with frequently recurring features within related cancer types [13] (Figure S2). This observation implies that particular CNA patterns may be specifically tolerated and/or contain elements which provide selective advantage during malignant transformation and disease progression.

Earlier research has described amplification and deletion hotspots among multiple cancer types [14, 15, 16, 17]. CNA-derived gene discovery can complement the knowledge of functional landscape during oncogenesis and pinpoint new genes previously unknown from point mutation analysis [18]. In particular, an integrative multi-cancer analysis for CNA-exerted susceptibility discovery can increase the statistical power to extract disease-relevant genes and delineate their functional impact across cancer types. In recent years, work from several data curation projects and international research consortia has led to an improved availability for generally compatible, genome-wide CNA profiling data with associated information and thereby enabled the development and benchmarking of integrative approaches. Particularly, the Progenetix CNA database has gathered 115 357 samples across 788 cancer types from published studies and cohorts, including the CNA data from 11 090 patients of 182 cancer types from the Cancer Genome Atlas (TCGA) Project[19, 20, 21].

In the last decade, GISTIC has been widely used to assess the significance of individual genomic regions in CNA data sets from individual genomic platforms [15, 22]. It uses a semiparametric permutation to calculate a score for each probe based on both amplitude and frequency and identifies regions significant for amplification and deletion. However, beyond the probe-level and region-level significance discovery, it does not offer a statistical test for gene-wise significance which would allow cross-study comparisons.

Here we describe an approach to evaluate gene-wise significance in CND which utilizes the non-random features in gene locus disruption. We verify our model by comparing the identified genes with known cancer driver gene sets as well as genes with reduced expression in the respective cancer types. Additionally, we corroborate the identified genes in terms of their biological impact with pathway analysis and cancer type clustering.

## 2. Results

We used data from three independent data sources depending on sample availability: 13 cancer types from the arrayMap collection, which represents a subset of the Progenetix database with available probe-specific genomic array data [23]; 12 cancer types from the TCGA project processed on genome-wide SNP6 arrays [19]; as well as 4 cancer types from cBioPortal database derived from whole exome sequencing (WES) experiments [24] (Table 1). Among these, 9 cancer types were represented by more than one source allowing comparison and benchmarking for source or technology related biases (Table 1). Their genome-wide CNA landscape differed among cancer types while remained comparable between data sources (Figure S3).

**Table 1:**
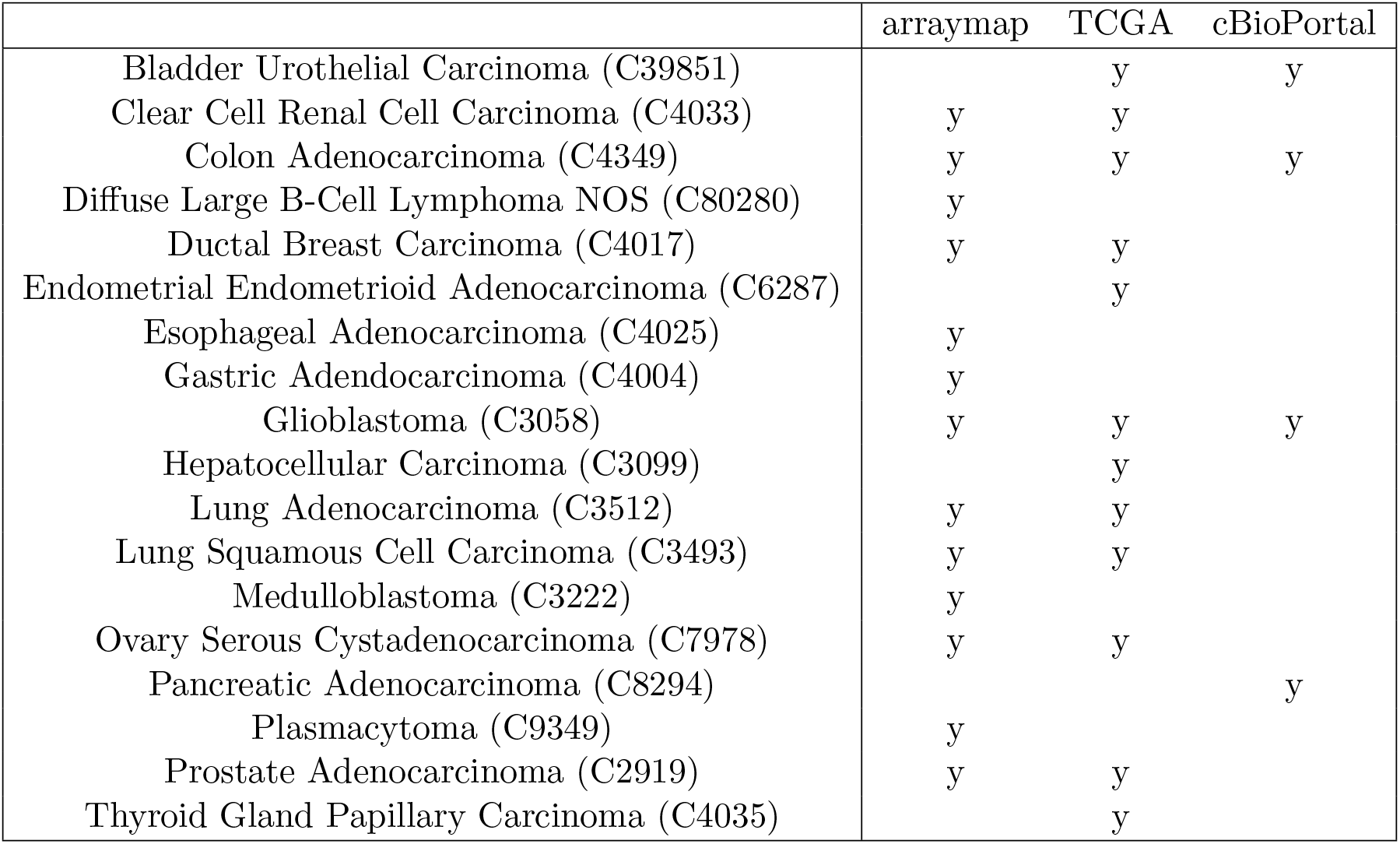
Analyzed cancer types across data sources

### 2.1. Model design

We observed non-random features in CND which can be harnessed to characterize their underlying mechanisms. First, we noted that CND segment size tend to be reduced in gene-rich regions. We overlaid the CND segments from all the datasets included in the analysis 1 and computed the number of genes hit by the collective segment profile normalized by segment length in units of 100kb, multiplied by the number of samples containing the segment. For simplicity, we refer to this value as gene hit frequency (GHF) from here. GHF decayed with increasing CND size (Figure 1). In the segment sizes below 5Mbp GHF decreased from 49.20 (bin 0, 0 - 100 kb) to 3.55 (bin 49, 4.9Mbp - 5Mbp) (Figure 1 Left). We noted that the individual GHF values in the first two bins’ (0 - 200 kb) upper quantile were above the axis limit and the individual GHF values spanned 5 orders of magnitude. Subsequently, we plotted the GHF for the first 5 bins in log scale (Figure 1 Right). Visibile on both scales, the individual points formed curves of discrete gene hit number (1, 2, 3…) divided by segment size (x-axis), while the zero-hit curve was not visible on the log-scale. We observed GHF with higher variability and higher median in the lower range of segment size and more consistently lower GHF as segment size increased, with a Spearman correlation coefficient at −0.11 and p-val 2.225 × 10^-308^ (python’s minimal float value). Overall, the GHF decay indicated that CND favored targeting specific genes and long un-targeted CND in gene-rich regions were selected against.

**Figure 1:**
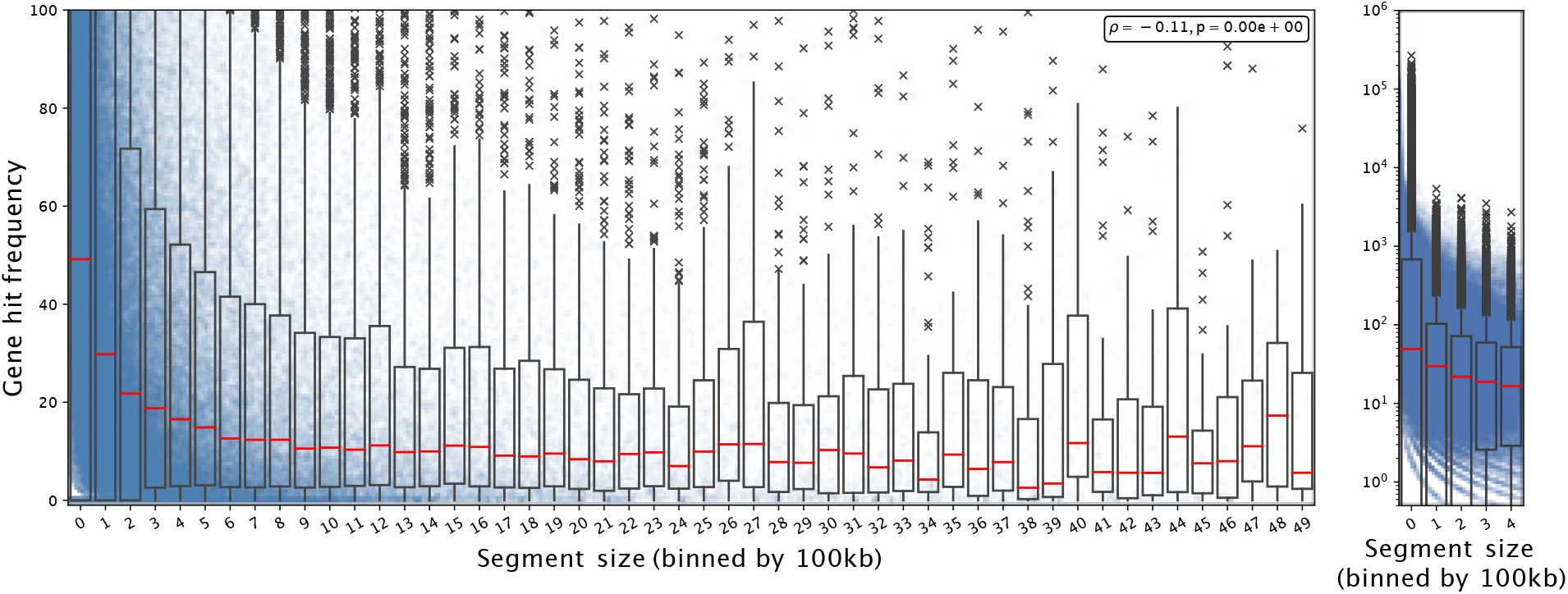
Gene hit frequency decreases as CND segment size increases. Each dot shows an individual gene deletion event in a segment falling into the bin size range and the boxplot shows the summary statistics of the bin including its 25, 50(red line; median), 75 percentiles. Left: The segment size is shown from 0 to 5MB; the gene hit frequency is limited up to 100 at a linear scale to show the decay with the bins. Right: for segment size between 0 and 500kb with log scale on y axis.

Further, we showed that CND tended to recur and to locate within the genomic range of cancer-related genes. We tested the recurrence with CND segment endpoints in expert curated driver genes from COSMIC [25] against the rest of the genome. The localization of breakpoints in driver gene sets is highly over-represented in all 29 analyses, with one-sided Fisher exact p value in the range of 0.038 to <2.225 × 10^-308^. After correcting for gene length, breakpoints were still over-represented in the driver set in 27 analyses, with one-sided Fisher exact p value in the range of 4.3817 × 10^-8^ to <2.225 ×10^-308^. For Diffuse Large B-Cell Lymphoma, NOS (C80280; dataset from arrayMap) the p value equaled 0.51 and for medulloblastoma (C3222; arrayMap) it equaled 1.

Based on these initial observations, we then designed a model to capture these nonrandom CND features as illustrated in Figure 2. In each cancer type, we aggregated all CND segments and created new “collective segments” in a reference genome track. We calculated a gene score for all the genes giving weight to the sample size on the segments covering the gene while penalizing the length of segments to account for size-related unspecific deletion.

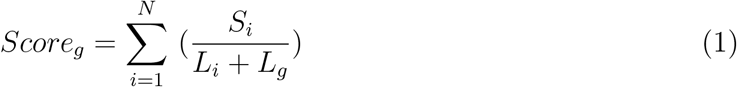

**Figure 2:**
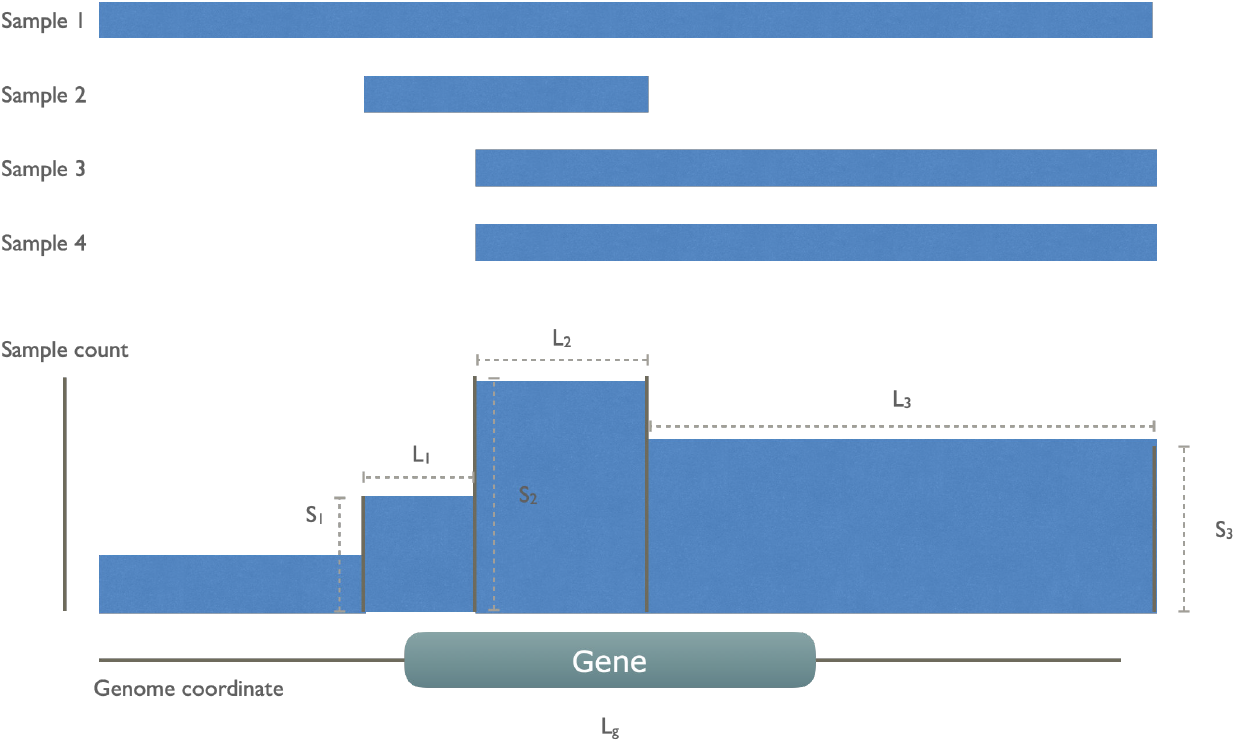
An illustration of gene score calculation. The gene score was defined to reward the high number of sample recurrence and penalize the length of segments and genes. In this example, there are four cancer samples, all of which have a whole or partial deletion on the indicated gene *g*. These CN segments are collapsed to a collective track, leaving 4 collective segments of which only 3 of them overlap with the gene. The gene score sums over these 3 segments (*i*) with count of involved sample *S_i_* divided by the sum of segment length *L_i_* and the common gene length *L_g_*.

For each gene *g*, a score is defined by summing up all overlapping deletion segments (N). For each segment *i*, the division of sample count *S_i_* and the sum of segment length *L_i_* and gene length *L_g_*.

The positions of collective CND segments were shuffled within the same chromosome. The gene scores were calculated for each shuffling, to generate a background score distribution for each gene. The gene score on the real data was compared with the background distribution to calculate a empirical p value to denote the gene’s significance, which was subsequently adjusted with Benjamini-Yekutieli procedure. Significant genes from each analysis had the adjusted p value below 0.05.

### 2.2. Significance across multiple cancer types

For the 29 datasets included in this study, we first assessed the breakpoint density in the gene-dense and gene-poor regions within each analysis. While genome-wide SNP array derived datasets from TCGA and arrayMap sources showed similar density, the WES-derived data from cBioPortal were biased against gene-poor regions, which causes inflation of gene significance level, making it not comparable with the array-derived data where probes are approximately equally distributed across the whole genome (Figure S4). Therefore, the WES data-based results were not used to derive a gene set by a significance cutoff but only for cancer type clustering with genome-wide significance p-values in Section 2.5.

In the 17 cancer types, the number of genes tested as significant with the segmental breakpoint model in each analysis ranged from 19 to 169 (Table S1). We used the three cancer-driving gene sets as the “gold standard” to test enrichment of identified genes in these sets [26, 27, 25]. 144 genes are included by all gene sets and 134 genes in at least two sets, while 784 genes are exclusively found in one set (Venn diagram in Figure S5; Gene set details in Appendix 3). In 20 out of 25 analyses, the identifed genes were enriched in the Bailey set (16/25 for Dietlein set and 18/25 for CGC set respectively; Table S2). Specifically, RB1 was found significant in 15 out of 25 analyses, followed by PTEN, CDKN2A, PTPRD, SMAD2, NRG1, JAK2, FHIT, DLC1, SMAD4, MAP2K4, RET, LRP1B, BRCA2, EPHA7, MLLT3, KANSL1, CTNNB1, APC, FGFR1, NCOR1, FLCN. Their functions spanned PI3K/Akt pathway, cell cycle regulation, Wnt signaling and chromatin histone modification pathways. The cross-study significant genes showed local clusters of significance, particularly in chromosome 8p, 9p and 17p (Figure 3), while a few others e.g. RB1, FHIT and PDE4D, appeared as singletons.

**Figure 3:**
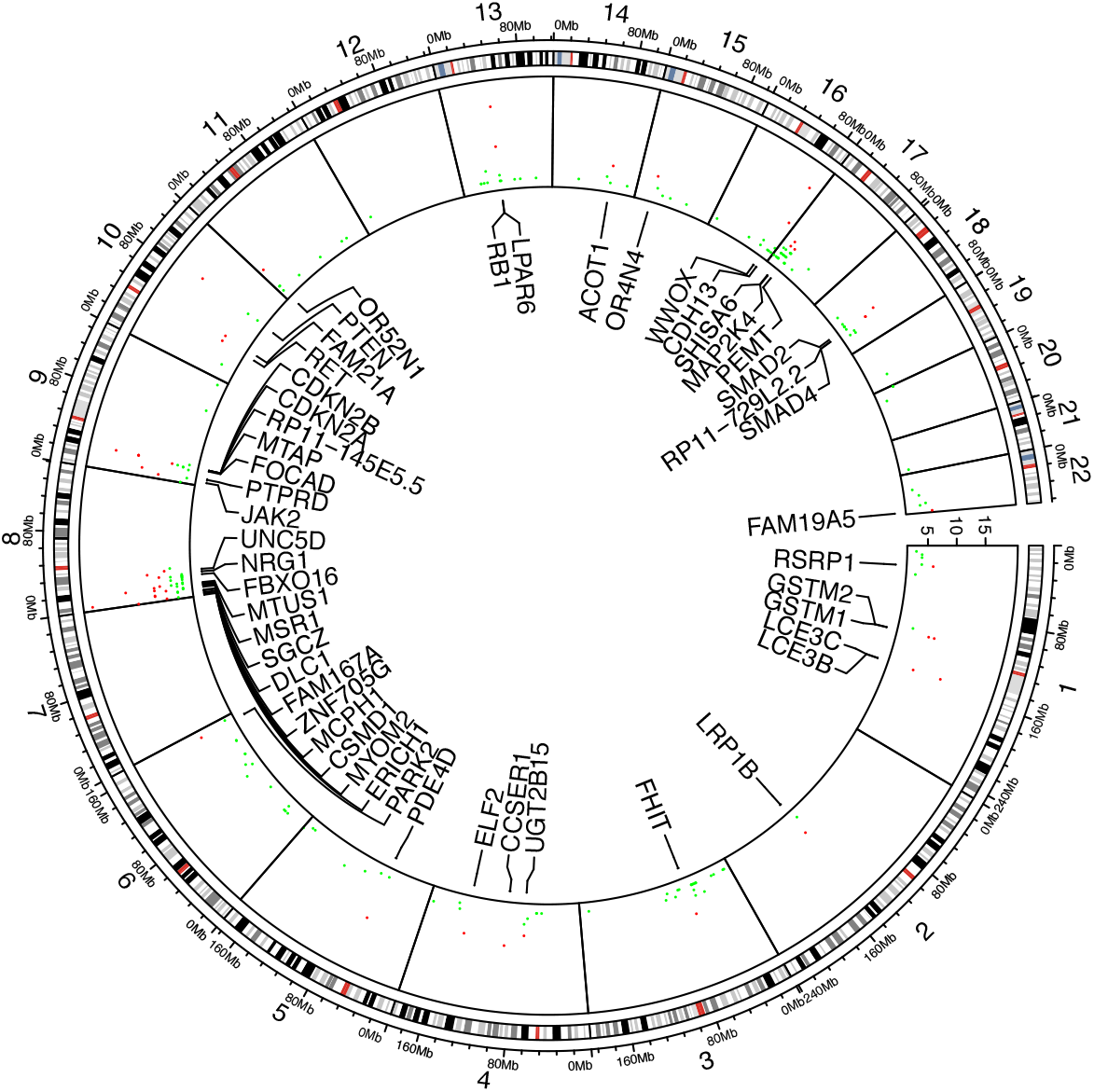
Significant genes shared among analyses. 49 genes found significant in more than 5 analyses are marked red and labeled accordingly.

### 2.3. Differential expression against normal tissue

Since cancer CNA has been previously found to be strongly correlated with gene expression, for a majority of genes, we expect that genes significantly covered by CND should have a reduced expression level compared to the matched normal samples [28]. We tested this hypothesis with the RNA sequencing data of paired cancer and normal samples from TCGA. For 9 cancer types with available data, we determined a validation list of genes with significant mRNA expression reduction in tumor samples as compared to normal samples. We expect that the reduced dosage at DNA level may result in a cascade effect on downstream effectors with potentially magnified impact, and that other somatic variations unrelated to copy number alterations may additionally influence a wide range of expression values. In turn, the significant genes should demonstrate an evident reduction in expression but may not be among the most down-regulated by log fold change. Here, analyses for all the cancer types showed significant over-representation of identified CND-significant genes in the down-regulated genes from RNAseq data (one-sided Fisher exact p values range from 1.6 × 10^-6^ to 5 × 10^-21^), corroborating the LOF at the expression level.

### 2.4. Pathway enrichment analysis

We also evaluated the dependency of the analyzed cancer types on different functional pathways. The clustering of pathways by significant genes showed a universal enrichment in the pathways related to cancer (Figure 4; zoomed area of 48 pathways). Among these were “TGF-beta signaling”, “stem cell”, “viral infection”, receptor signaling pathways related to growth and apoptosis, “senescence”, “energy metabolism”and “cell adhesion”. With paired cancer types derived from different data sources, two cancer types out of nine - ovary serous cystadenocarcinoma (C7978) and glioblastoma (C3058) - were clustered together. Furthermore, a list of 29 canonical pathways from Supplementary Table 5 of [1] were used as hallmarks in cancer development. For these pathways, clustering was performed without standardization to compare the influence among cancer types as well as between multiple resources (Figure S1). It was conspicuous that none of the analyses was enriched in mismatch repair pathway(MMR). MMR is common for familial cancers, including hereditary non-polyposis colorectal cancer (HNPCC) or Lynch syndrome [29]. Microsatellite instability (MSI) caused by MMR defects is mutually exclusive with the chromosomal instability related to CNA [30], which is in accordance with our result. Three pathways, “TGF-beta signaling”, “Cell cycle” and “p53 signaling”, were enriched in a majority of cancer types. Additionally, Ras signaling pathway was enriched in prostate adenocarcinoma from arrayMap dataset and GnRH signaling pathway was enriched in ovary serous carcinoma from TCGA dataset.

**Figure 4:**
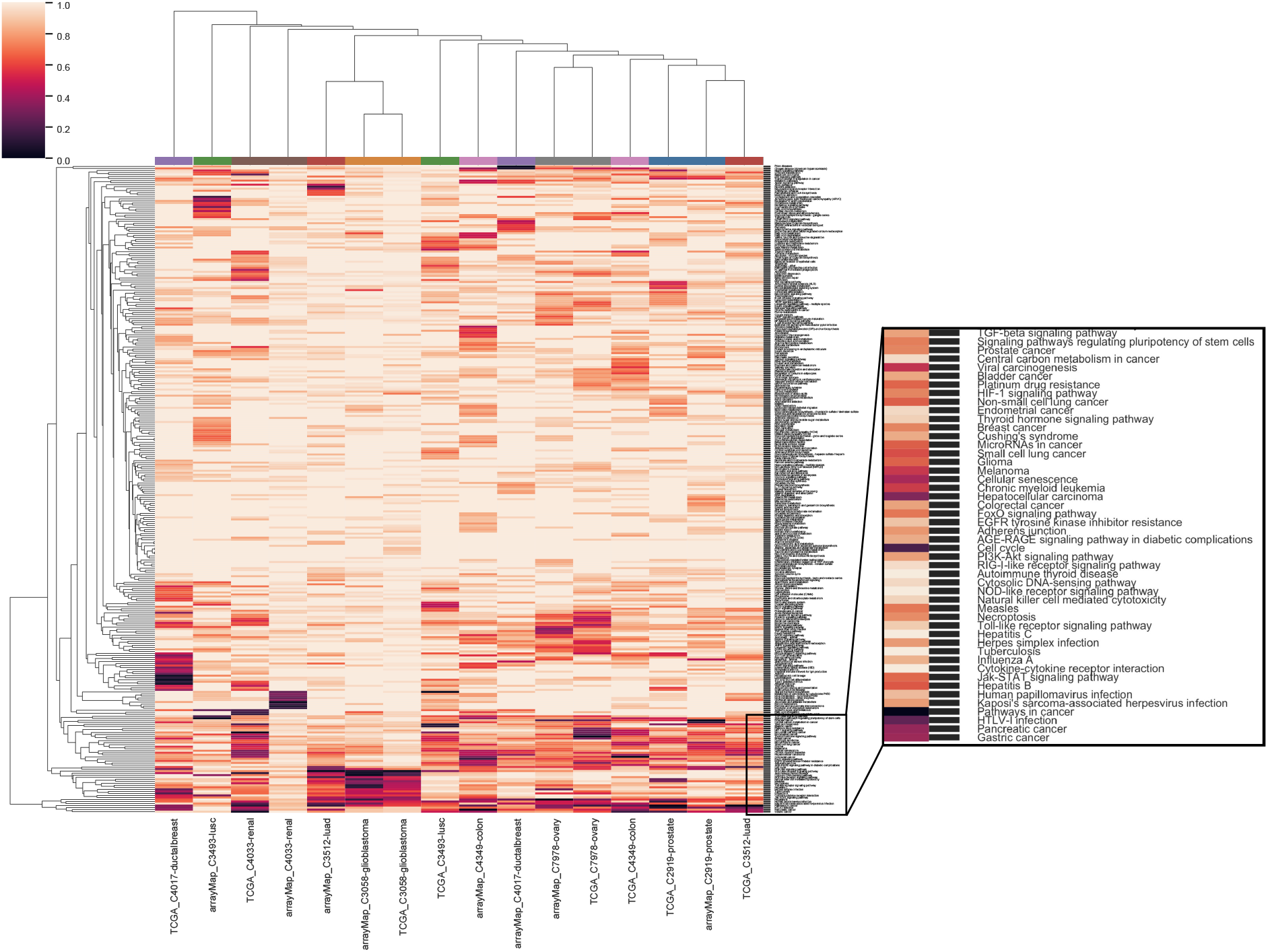
Clustering of genome-wide significance value of paired 8 cancer types with all 326 KEGG pathways. The same color coding of analyses on the top of the heat map indicate the same cancer type.

We listed the top 10 pathways by enrichment significance in the 25 analyses (Table S3). “Pathways in cancer” appears in 11 analyses. Different types of viral infection appear in 12 analyses while different types of drug metabolism appear in 9 analyses. “Metabolism of xenobiotics by cytochrome P450”, “Cell cycle” and “Cellular senescence” appear in 7 analyses. “Chemical carcinogenesis” appears in 6 analyses.

### 2.5. Cancer type clustering

So far, we used FDR cutoff to identify significant genes within an analysis. However, among the genes not passing the threshold, their p-values entailed information and constituted a pattern which could distinguish between cancer types. To test this hypothesis, we used the genome-wide significance scores to assess similarities between cancer types. We used uniform manifold approximation and projection (UMAP) to reduce the gene significance p-values to two dimensions and indicated identical cancer types from different data sources with the same color (Figure 5). All matched cancer types were represented in a neighborhood, except for prostate adenocarcinoma (C2919). The ability to differentiate between cancer types substantiated that each gene’s p-value from the method contained additional information about the analyzed cancer type.

**Figure 5:**
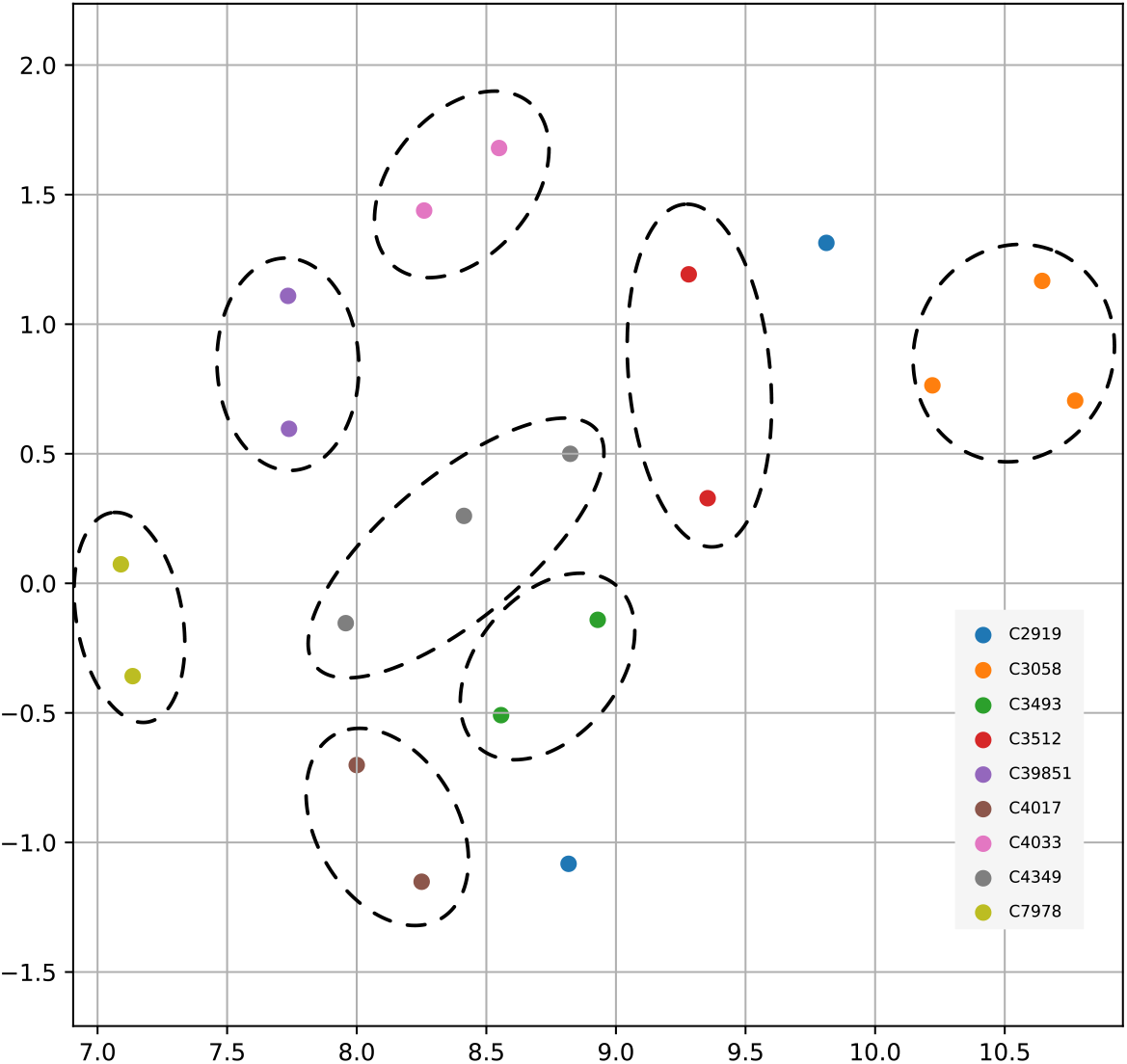
UMAP projection of genome-wide significance scores among 9 paired cancer types from multiple data sources. In order of appearance the NCIt codes in the figure legend represent - C2919: Prostate Adenocarcinoma; C3058: Glioblastoma; C3493: Lung Squamous Cell Carcinoma; C3512: Lung Adenocarcinoma; C39851: Bladder Urothelial Carcinoma; C4017: Ductal Breast Carcinoma; C4033: Clear Cell Renal Cell Carcinoma; C4349: Colon Adenocarcinoma; C7978: Ovarian Serous Cystadenocarcinoma

## 3. Discussion

We have proposed a data-driven method based on the frequency of gene disruption by segmental breakpoints to discern non-random CND at a gene level and generated a list of genes significantly affected by CND as well as genome-wide significance scores. From 29 independent runs on 18 cancer types, we have benchmarked the cancer-driving effects of the resulting genes through enrichment analysis for three independent “driver” gene sets and found significant enrichment all three sets. In addition, the genes show significant enrichment for cancer-related pathways and reduction in mRNA expression. Using the genome-wide significance scores we have clustered the analyses from multiple independent data resources and showed moderate separation between cancer types.

Apart from providing gene-wise significance measure, this method has several advantages, including the preservation of gene neighborhood information, robustness with respect to global CNA content and variation from individual samples. First, the shuffling preserves the genomic context and gene neighborhood structure. Consider two genes A and B in close proximity. If they are within the same CND segment (co-segregation, equal disease relevance), the randomization would give them same background sample count. The gene length difference is also reflected in the background rate and they have the same effect on the significance score. In contrast, if gene A is more disease-relevant, there would be more CND segments overlapping with gene A, resulting in smaller collective segments for gene A score calculation. With the similar background rate due to location proximity, gene A will hence a higher score compared to gene B. In addition, the method accounts for the variation in overall CNA content. Specifically, if a set of samples mostly consists of profiles with an overall low amount of CNA, the probability of CNA at each given region remains and will not affect the outcome of the analysis. Finally, it is robust for the variation introduced from individual samples. Individual samples’ segmentation results can differ based on baseline setting - shift between adjacent CN states [31], but such shifting-derived error does not substantially affect the final outcome of the method, as a single sample count has little influence on the modelled aggregation of the sample collection

On the other hand, this method requires a dataset selection that compromises between sample size and cancer type specificity (for a representative and comparable CND profile), as a mixture of cancer subtypes with highly variable CNA landscape introduces breakpoint bias, and conversely a small sample size and few breakpoints result in low statistical power. Also, it is expected that in cases of chromothripsis-like events (CTLP; focal, extreme hypersegmentation) [32, 33] the significance score of local genes can be increased and the significance of other genes on the same chromosome can be reduced due to the rise in the overall CNA rate. Such samples should be pre-filtered with the criteria summarized in [34] and investigated carefully on a case-by-case level.

Genes with significant scores across multiple analyses include established tumor suppressor genes that control cell cycle and regulate proliferation and programmed cell death [35]. RB1 is implicated in multiple processes, including cell cycle, stress responses and apoptosis[36]. FOXO and PTEN are key regulators in phosphatidylinositol 3–kinase (PI3K) pathway which reacts to growth signals[37, 38]. CDKN2A(p16), CDKN2B(p15) at 9p21 controls S-phase checkpoint and their deletions have been reported in multiple cancers [39, 40], SMAD2/4 are involved in TGF-beta signaling pathway [41, 42]. The list also includes genes with sporadic or ambiguous oncogenic attribution. FGFR1 has been reported to have both oncogenic and tumor suppressive potential [43] and the tyrosine kinase RET has long been established as a classic proto-oncogene but was found to act as a TSG in colorectal carcinomas [44, 45].

As many high-level regulators have alternative cancer-promoting roles depending on the cellular context, the evidence of their selective deletion in a multi-cancer analysis provides additional support for their relevance in oncogenic processes. Our results also point out genes with emerging cancer-related roles outside of classical cancer pathways such as GSTM1/GSTT1 in xenobiotic metabolism [46], ELF2 as an ETS transcription factor regulating various biological pathways [47] or DLC1 as a Rho-GTPase activating protein regulating *RhoA* pathway in hepatocellular carcinoma [48]. In addition, we have identified large genes residing in common fragile sites to be significantly affected by deletions and contributing to cancer development, including CSMD1, WWOX and FHIT [49].

Through the pathway analysis, we have observed prevalent enrichment in cancer-related pathways as well as hallmark mechanisms for cancer progression. In summary, the TGF-beta signaling pathway, cell cycle regulation and p53 signaling pathways emerged as the most frequently affected among all cancer types. Prostate, ovary and ductal breast adenocarcinoma samples were enriched for a majority of hallmark pathways, confirming their prominent dependence on CNA compared to discrete mutational events, as previously established[50].

The genomic CNA patterns as in Figure S3 have been trained to distinguish cancer types through binned genome-wide CNA status [14], but the predictive signatures to the level of chromosomal regions are not readily interpretable for biological functions. On the contrary, gene-level scores for cancer type clustering in our analysis shows clear but limited separation. This demonstrates that the cancer-type-specific CN signature is captured in the genome-wide scores. The limited distinction may be related to the vast amount of non-coding areas not included [51]. Indeed, as an example in the local context of CDKN2A/B, a long non-coding RNA ANRIL is responsible for the transcriptional regulation, miRNA interaction, which modulates proliferation, senescence, motility and inflammation[52]. Additionally, heterogeneity within the sample set, non-CNA driven cancer samples as well as shared significance of core CNA genes may have overshadowed the less impactful cancer-type-specific genes for the cluster separation.

In this article, we have developed a method to extract non-random significance from copy number deletion in cancer which exploits the specific functional implications of genomic deletion or disruption events. While due to the differences in genomic architecture and functional mechanisms of copy number gains this method is not suited to deliver a ”universal CNA model” in principle the method could be adapted to discern non-random copy gain significance. Namely, cancer-promoting genes could be expected to show under-represented disruption by either gain or loss copy endpoints as well as over-representation of endpoints in close proximity to gene start and end which however should have overall results distinct from the observed CND statistics.

In summary, we provide a general framework for integrative analysis on copy number deletion. It has confirmed well-known tumor suppressor genes as well as identified genes with incomplete characterization of their mode of action, suggesting the value in novel discovery and promoting further research into less studied genes. With the growing collection in high-quality CNA data, this method can be expanded to rare cancers which will potentiate discovery of novel cancer susceptibilities and dependencies and complement the overall understanding of malignancy development. Confirmed by the functional characterization of the known coding genes, this tool might be extended to the non-coding area and provide a better overview of the CNA functional landscape.

## 4. Experimental procedures

### 4.1. Data availability

CNA data has been accessed from three different sources - arrayMap, TCGA, cBioPortal - which had been integrated into the Progenetix database (Table 1)[23, 19, 24, 21]. CN data and curated biosample metadata are freely accessible through progenetix.org over the GA4GH Beacon protocol in JSON format compatible to the Beacon v2 data model as well as tab-delimited text file format [53, 54, 55].

For differential expression analysis, transcriptomics data in raw HT-Seq counts was accessed for respective TCGA projects from GDC Data Portal. Paired RNAseq data was available for 11 cancer types: prostate adenocarcinoma (C2919), hepatocellular carcinoma (C3099), lung adenocarcinoma (C3512), ductal breast carcinoma (C4017), thyroid gland papillary carcinoma (C4035), endometrial endometrioid adenocarcinoma (C6287), lung squamous cell carcinoma (C3493), bladder urothelial carcinoma (C39851), clear cell renal cell carcinoma (C4033), colon adenocarcinoma (C4349), ovary serous cystadenocarcinoma (C7978).

### 4.2. Differential expression analysis

We used R-package EdgeR for differential expression analysis between tumor and normal groups [56]. For each cancer type, gene-wise counts from paired tumor — normal samples were used. A TMM normalization was performed to calibrate for the library size (total counts) per sample. Negative binomial model was used to estimate the common and genewise dispersion parameters. A gene-wise general linear model was fit by the paired design and the differentially expressed genes were determined by likelihood ratio test.

### 4.3. Pathway analysis

All 326 KEGG pathways [57] were used to determine the enrichment in each pathway with a one-sided Fisher exact test with the contingency table of significant genes - genes with a significance score on one axis and genes in/not in pathway on the other axis. For clustering analysis including all pathways, log10 of Fisher exact p values were calculated and standardized to 0-1 scale, i.e. 0 with lowest p. Hierarchical clustering with Euclidean distance and average linkage method was performed on both cancer types and pathways. For the clustering analysis of the 29 canonical cancer pathways, the original Fisher exact p value was used. Hierarchical clustering with Euclidean distance and average linkage method was performed on pathways only.

## 5. Acknowledgement

We would like to thank the scientific input from members of the Zurich Seminars in Bioinformatics as well as the Theoretical Cytogenetics and Oncogenomics group at University of Zurich for continuous work on data collection and curation for the Progenetix database.

## 6. Author contributions

QH conceived the project, performed analysis and wrote the manuscript. MB provided the CNA data assembly, gave insights about data analysis and clinical cancer biology and edited the manuscript.

## Appendix 1. Significance across cancer types

**Table S1:**
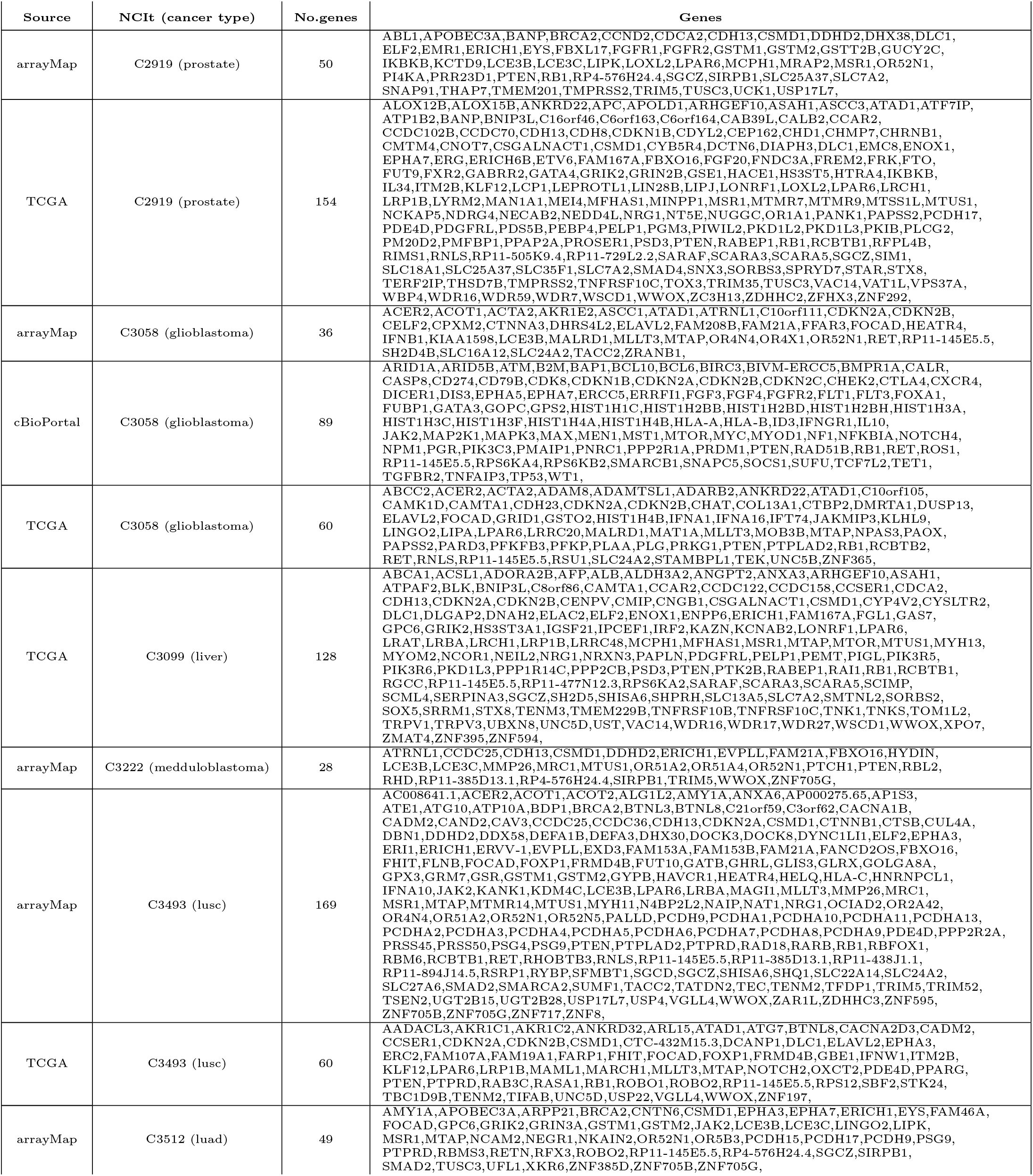
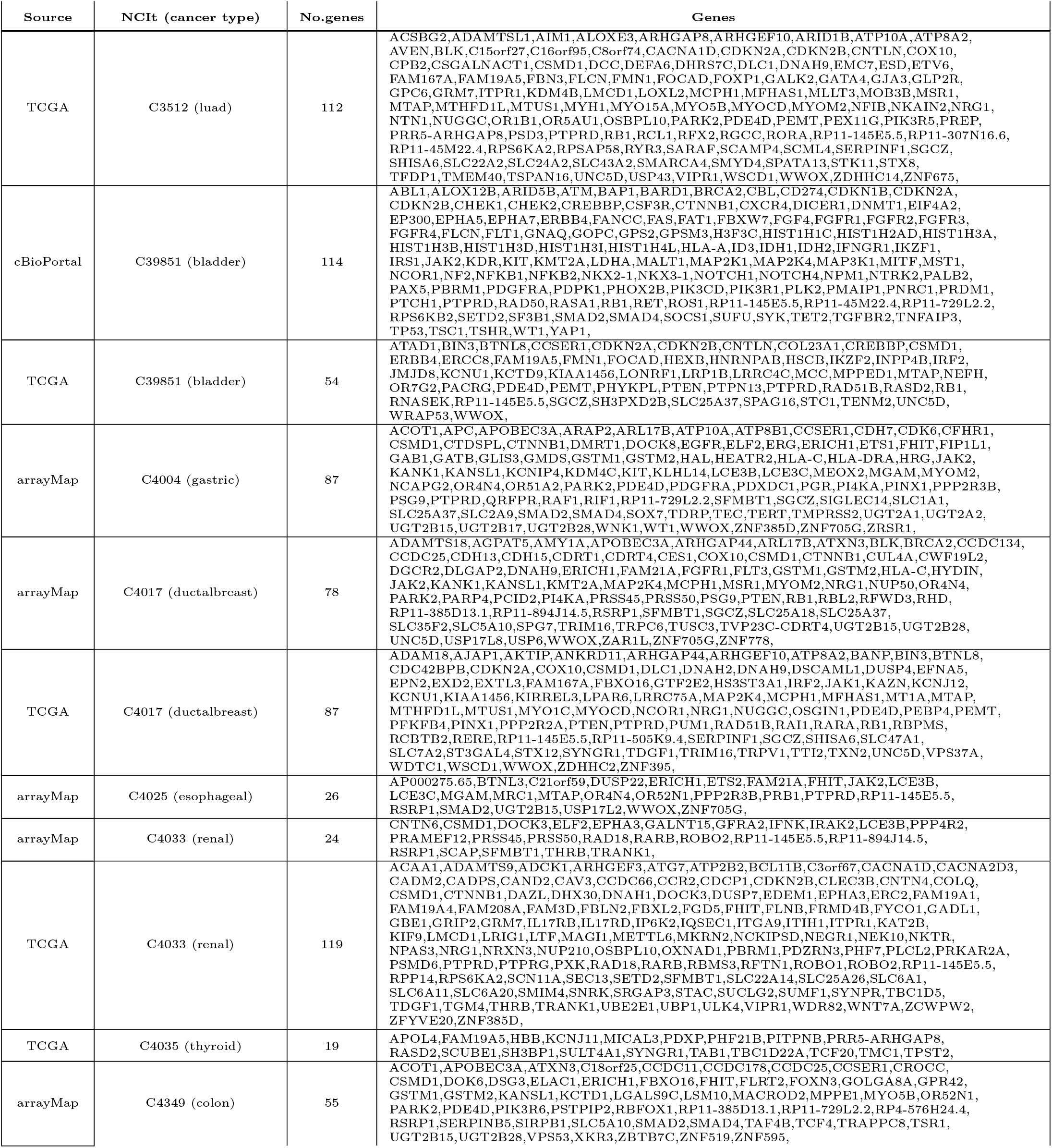
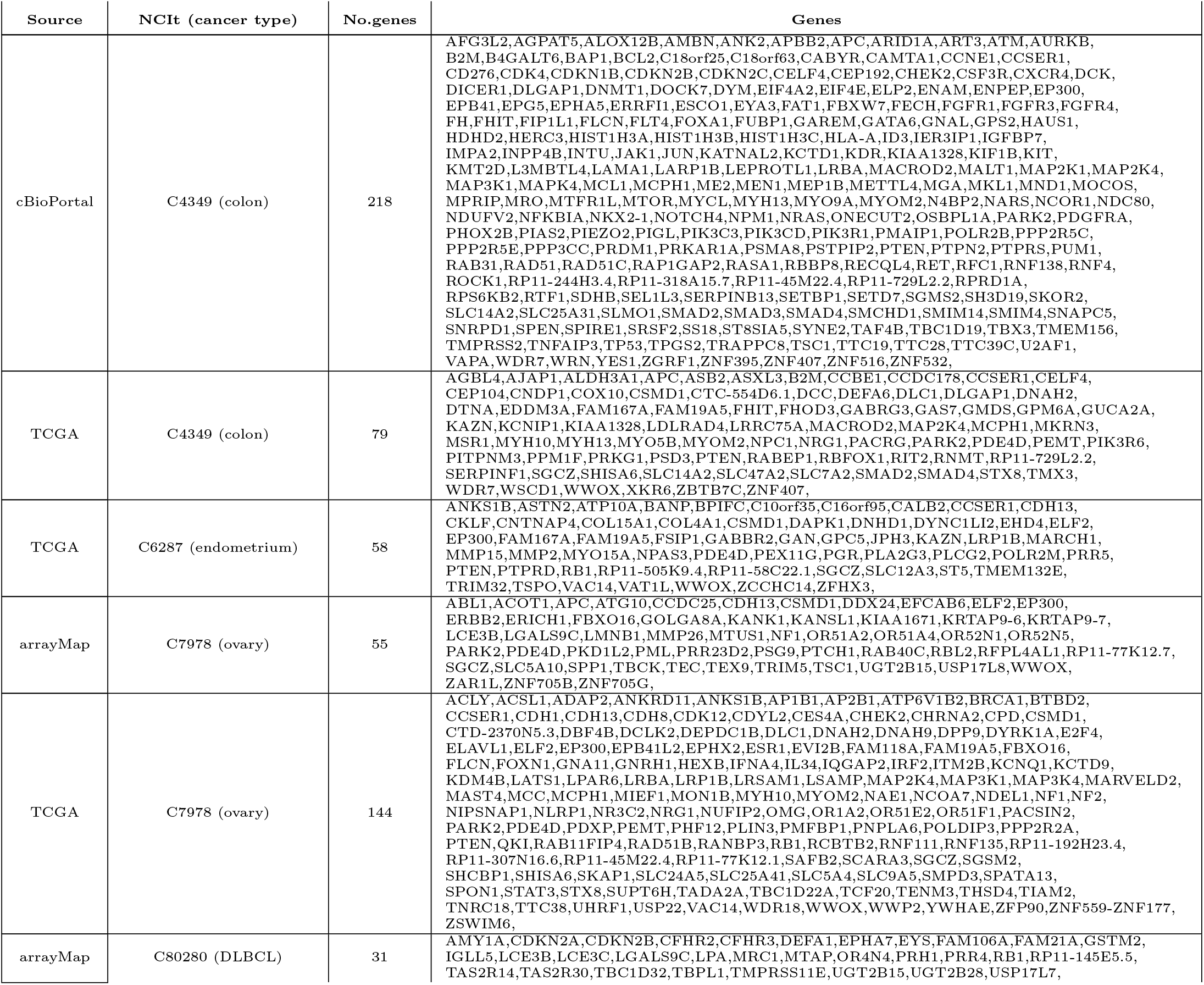
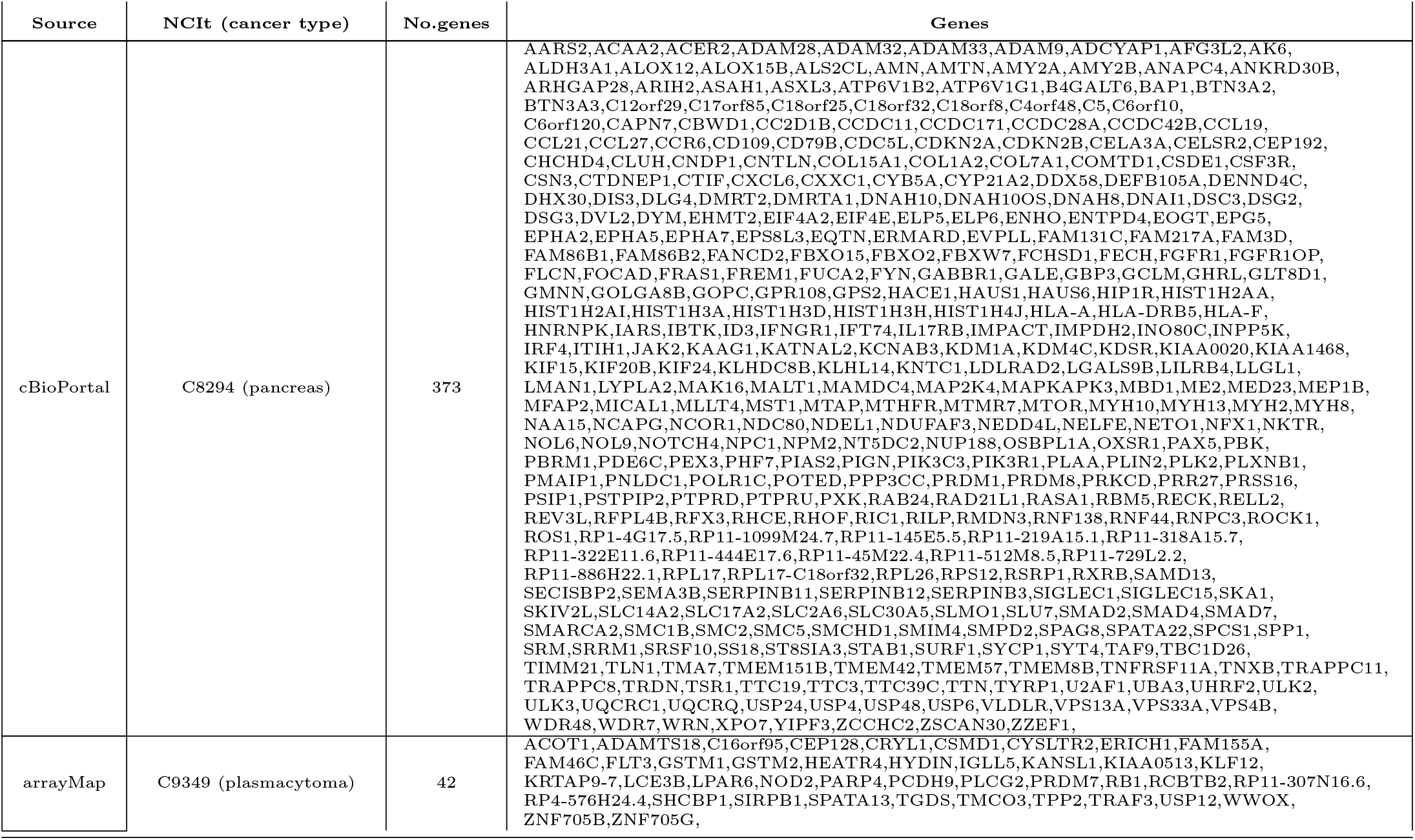
Significant genes in analyses

**Table S2:**
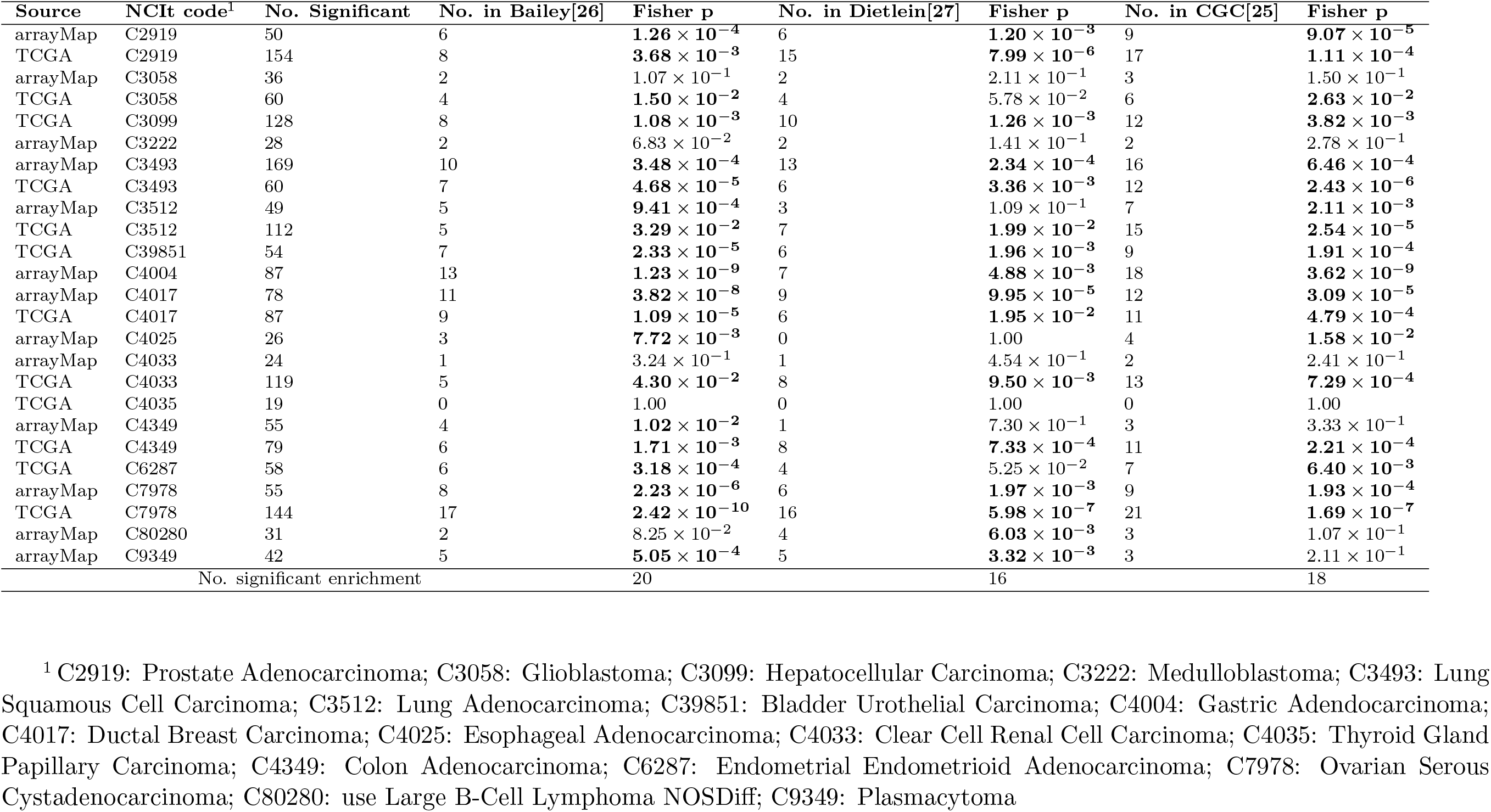
The number of significant genes belonging to the three cancer driving gene sets and the one-sided Fisher exact test p value for each analysis

## Appendix 2. Pathway enrichment and clustering

**Figure S1:**
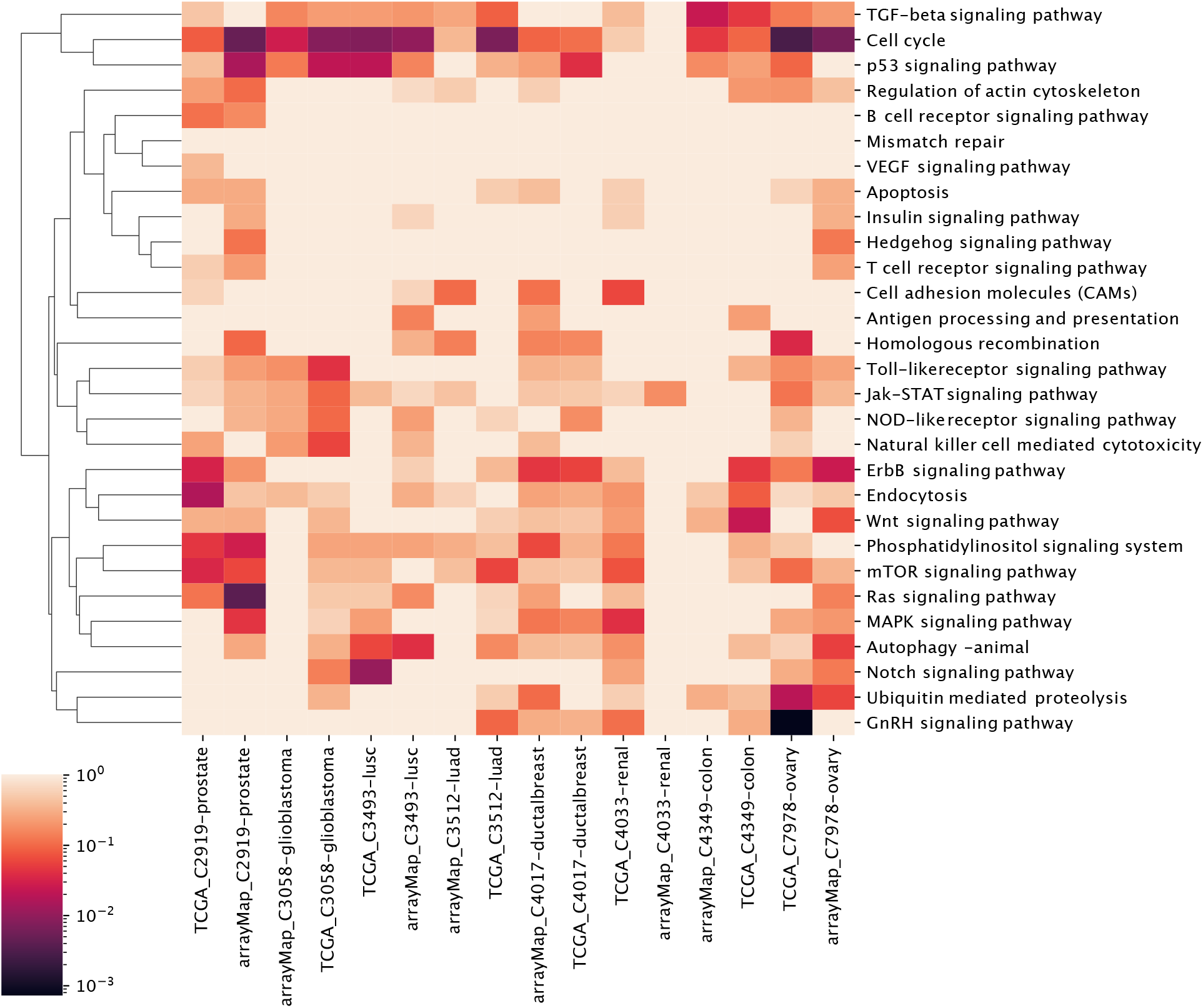
Clustering of genome-wide significance value of paired 8 cancer types with 29 cancer hallmark pathways.

**Table S3:**
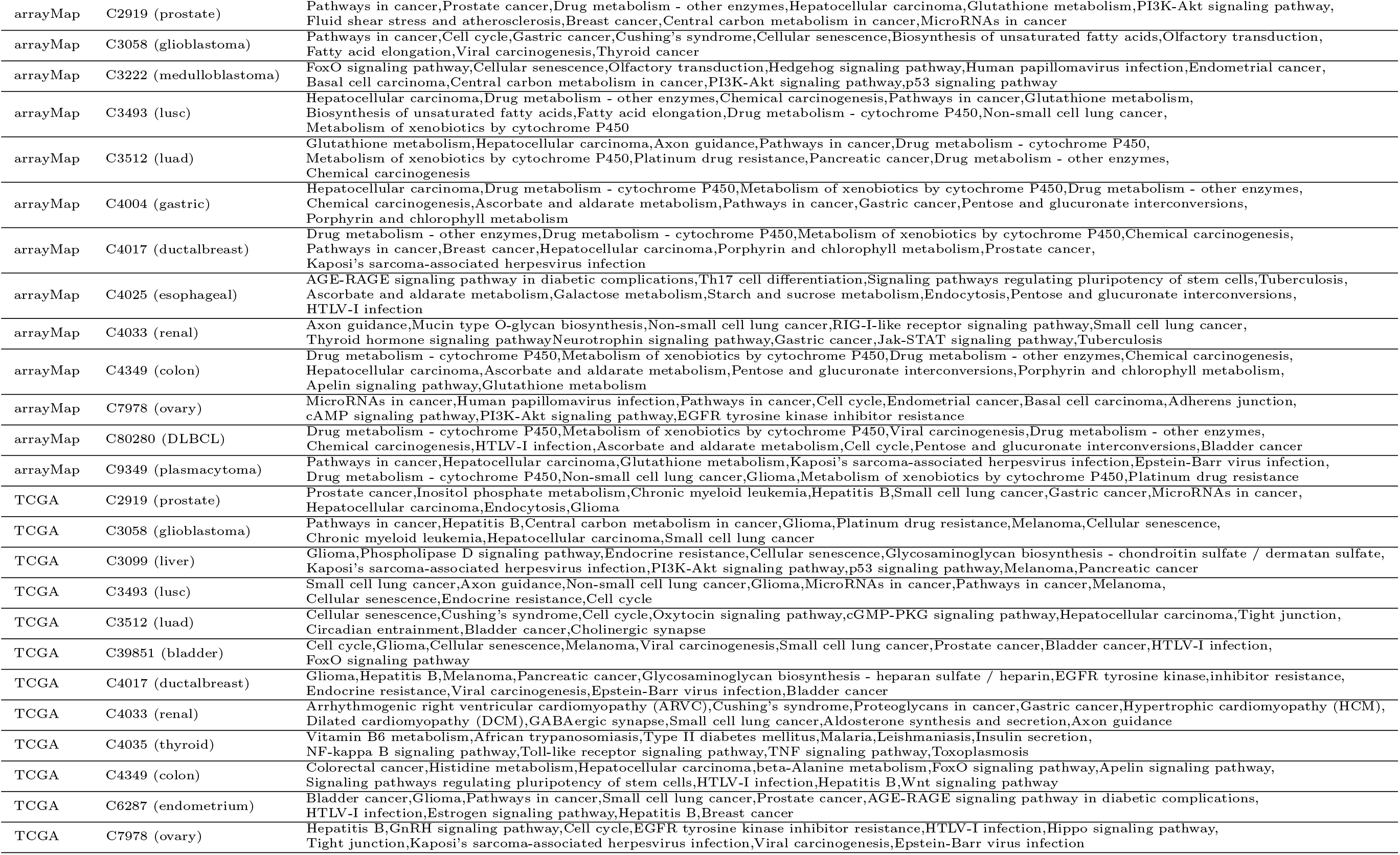
Top 10 pathways enriched by the significant genes of 25 analyses

## Appendix 3. Technical information of analyses and benchmarking sets

We have used GRCh38 (Ensembl Release 77) for the reference genome track and the gene CDS positions in the study.

Three independent driver sets include Bailey set [26] consisting of 299 genes from tumor exome analysis with experimental validation, the Dietlein set [27], including 461 genes from nucleotide context as well as the CGC set [25], including 724 genes from expert curation across multiple cancer types. The three gene sets share 144 consensus genes and the number of genes private to one set ranges from 73 to 481 (Figure S5).

**Figure S2:**
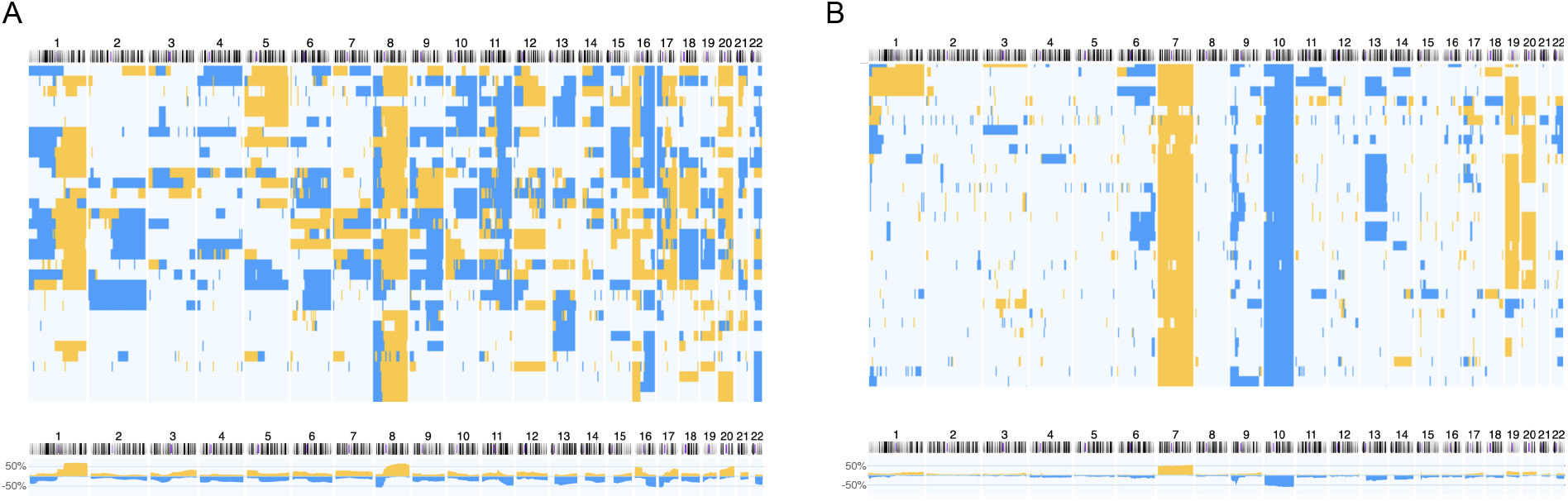
Genome-wide copy number landscape across multiple samples. A) Ductal Breast Carcinoma, NCIT:C4017; B) Glioblastoma, NCIT:C3058. In each cancer type, the top panel stacks 30 randomly selected individual samples’ CNA profile and the lower panel indicates the aggregated CNA landscape by percentage of samples exhibiting gain (yellow) and loss (blue) across the genome.

**Figure S3:**
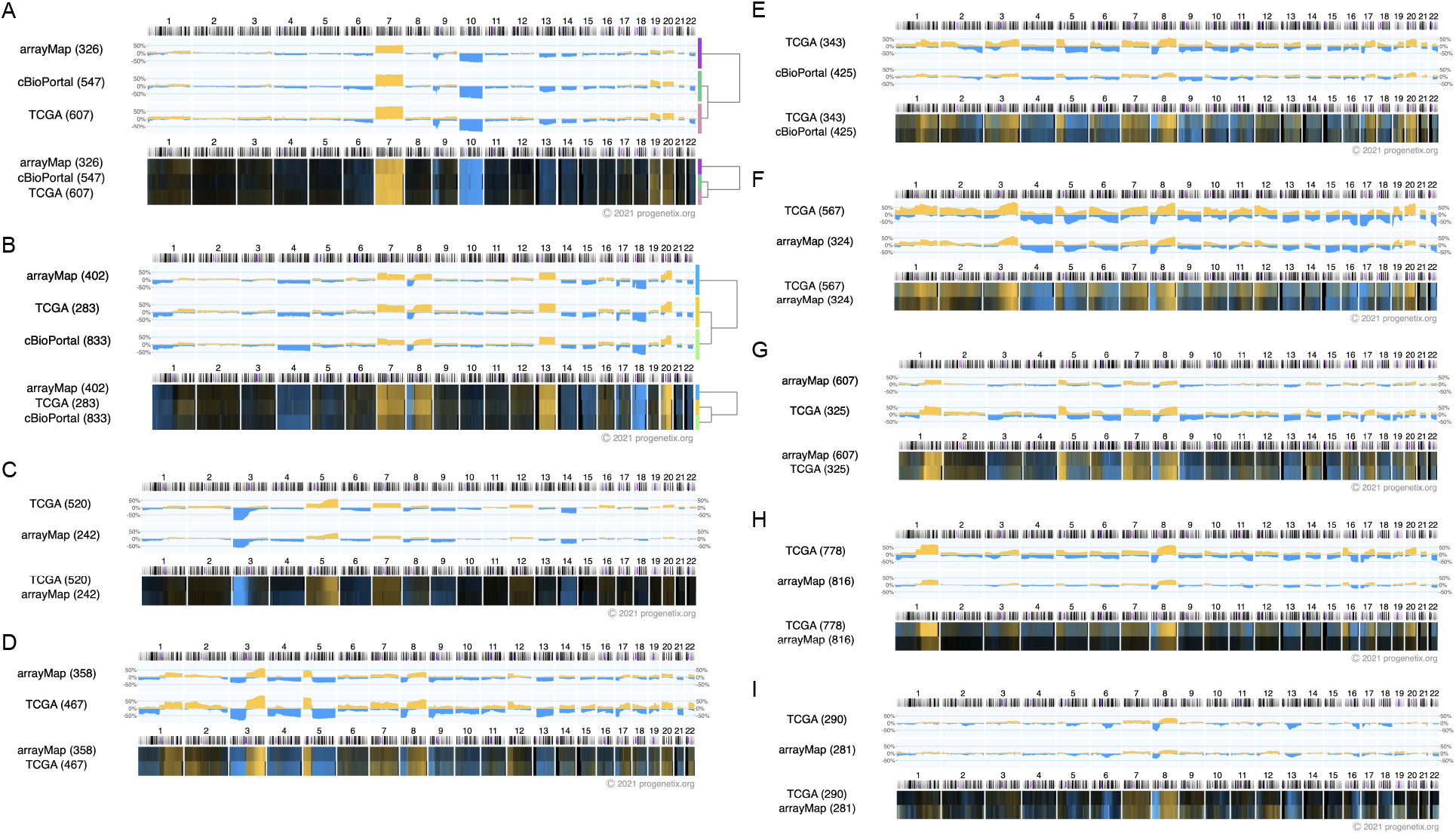
Genome-wide copy number landscape of nine cross-comparison cancer types. A) glioblastoma (C3058); B) colon adenocarcinoma (C4349); C) clear cell renal cell carcinoma (C4033); D) lung squamous carcinoma (C3493); E) bladder urothelial carcinoma (C39851); F) ovary serous cystadenocarcinoma (C7978); G) lung adenocarcinoma (C3512); H) ductal breast carcinoma (C4017); I) prostate adenocarcinoma (C2919).

**Figure S4:**
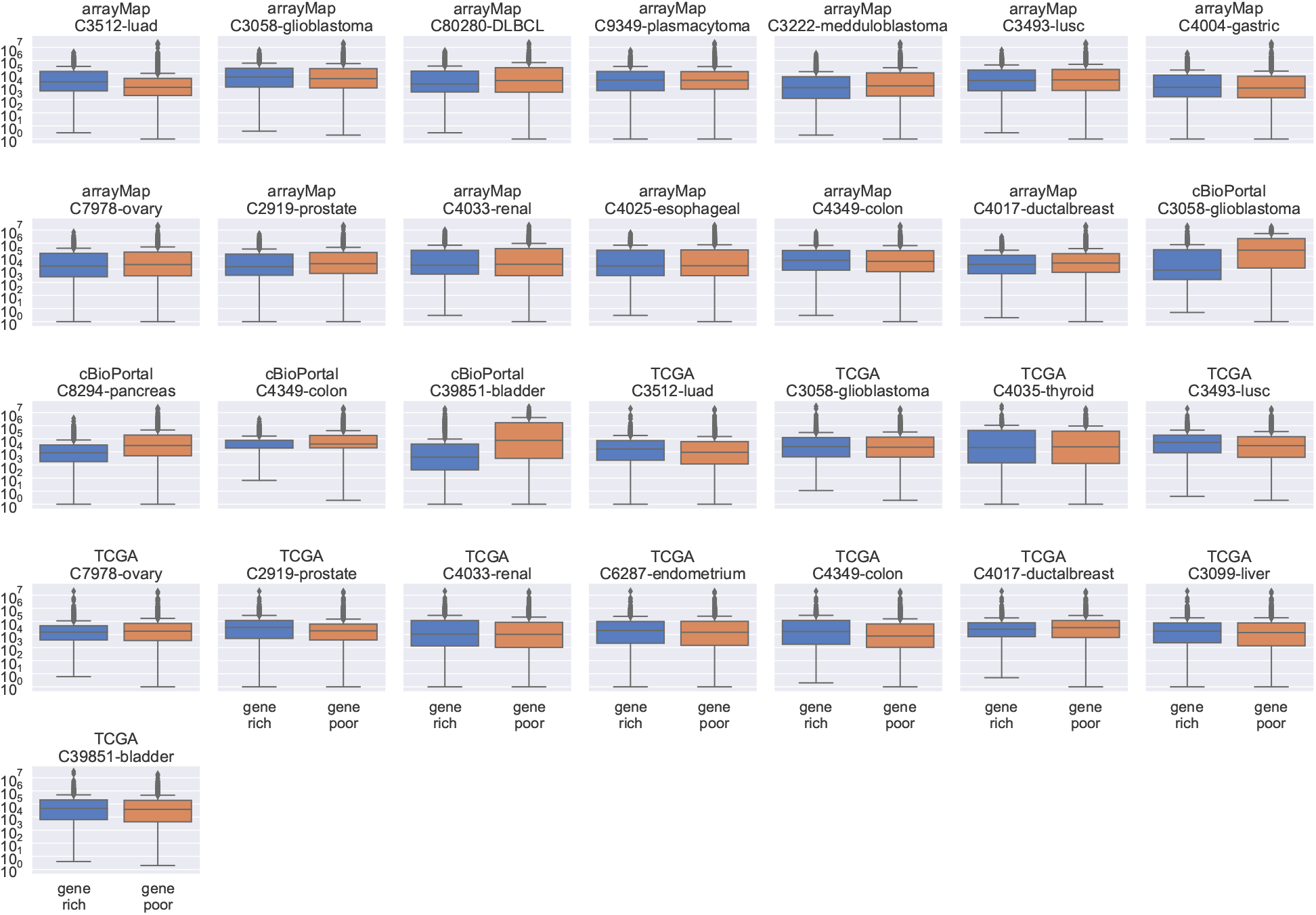
Number of segmental breakpoints in gene-rich and poor regions by analysis. Most anal-yses have a nearly equal breakpoint density regardless of gene density but the four WES-derived (cBioPortal) analyses show clear segment sparsity in gene-poor regions.

**Figure S5:**
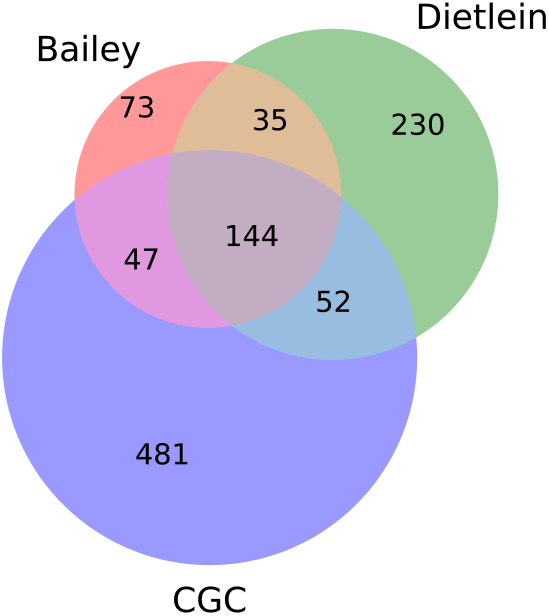
Overlap among the three cancer driving gene sets: Bailey, Dietlein and CGC [26, 27, 25].

